# Spatial Analysis of Neural Cell Proteomic Profiles following Ischemic Stroke in Mice using High-Plex Digital Spatial Profiling

**DOI:** 10.1101/2021.08.25.457708

**Authors:** Jessica M. Noll, Catherine J. Augello, Esra Kürüm, Liuliu Pan, Anna Pavenko, Andy Nam, Byron D. Ford

**Author notes:** Corresponding Author: Byron D. Ford Ph.D., University of California-Riverside School of Medicine, Division of Biomedical Sciences, 900 University Ave. Riverside, CA 92521.

## Abstract

Stroke is ranked as the fifth leading cause of death and the leading cause of adult disability. The progression of neuronal damage after stroke is recognized to be a complex integration of glia, neurons, and the surrounding extracellular matrix, therefore potential treatments must target the detrimental effects created by these interactions. In this study, we examined the spatial cellular and neuroinflammatory mechanisms occurring early after ischemic stroke utilizing Nanostring Digital Spatial Profiling (DSP) technology. Male C57bl/6 mice were subjected to photothrombotic middle cerebral artery occlusion (MCAO) and sacrificed at three-days post-ischemia. Spatial distinction of the ipsilateral hemisphere was studied according to the regions of interest: the ischemic core, peri-infarct tissues, and peri-infarct normal tissue (PiNT) in comparison to the contralateral hemisphere. We demonstrated that the ipsilateral hemisphere initiates distinct spatial regulatory proteomic profiles with DSP technology that can be identified consistently with the immunohistochemical markers, FJB, GFAP, and Iba-1. The core border profile demonstrated an induction of neuronal death, apoptosis, autophagy, immunoreactivity, and early degenerative proteins. Most notably, the core border resulted in a decrease of the neuronal proteins Map2 and NeuN, an increase in the autophagy proteins BAG3 and CTSD, an increase in the microglial and peripheral immune invasion proteins Iba1, CD45, CD11b, and CD39, and an increase in the neurodegenerative proteins BACE1, APP, αβ 1-42, ApoE, and hyperphosphorylated tau protein S-199. The peri-infarct region demonstrated increased astrocytic immunoreactivity, apoptotic, and neurodegenerative proteomic profile, with an increase in BAG3, GFAP, and hyperphosphorylated tau protein S-199. The PiNT region displayed minimal changes compared to the contralateral cortex with only an increase in GFAP. Overall, our data highlight the importance of identifying ischemic mechanisms in a spatial manner to understand the complex, dynamic interactions throughout ischemic progression and repair as well to introduce potential targets for successful therapeutic interventions.

## Introduction

Ischemic stroke is the leading cause of serious long-term disability and the 5^th^ leading cause of death in the United States (Benjamin 2017). Ischemic stroke represents approximately 80% of strokes and presents as the occlusion or blockage of an artery within the brain (Benjamin 2017). Currently, only tissue plasminogen activator (tPA) is approved for treatment of stroke, but has a limited time window for therapeutic use and is only administered to 3-5% of patients (Fisher et al. 2009, Bosetti et al. 2017). However, tPA treatment does not play a neuroprotective role, and is not ideal for prevention of secondary, long-term neurodegenerative damage. Additionally, as stroke progression is recognized to be a complex integration of glia, neurons, and the surrounding extracellular matrix, treatments should target the detrimental effects created by these interactions (Detante et al. 2014, Jayaraj et al. 2019, Jeong et al. 2013). Therefore, there is a strong need to understand the cellular and molecular mechanisms occurring early after ischemic stroke in a spatial manner in order to develop effective ischemic stroke treatments.

Ischemic stroke begins with loss of glucose and oxygen that triggers necrotic neuronal death (infarct), release of oxygen radicals, microvascular injury, and blood-brain barrier disruption in the ischemic core (Jayaraj et al. 2019). This then leads to secondary apoptotic, excitatory neuronal death in the surrounding ischemic penumbra with increases in neuroinflammation that can be prolonged for days after injury (Hossmann 2006, Brouns and De Deyn 2009). After ischemic damage is complete, the remaining, bordering region of reversibly damaged cells in an area of decreased neuronal density is defined as the peri-infarct tissue (Lipton 1999). The peri-infarct region exhibits a complex, dynamic interaction between immune and neuronal cells in both rodents and humans that critically determine late neuroprotection and repair processes after ischemia (Price et al. 2006, Cramer et al. 2006). Neuroinflammation after ischemia involves activation of microglia, astrocytes, and infiltration of peripheral leukocytes (Hallenbeck 1996, Kim et al. 2016, Pekny and Nilsson 2005, Kawabori and Yenari 2015). Resident microglia demonstrate reactivity with a change in morphology and infiltration into the core (Nowicka et al. 2008, Kawabori and Yenari 2015, Sanchez-Bezanilla et al. 2021). Astrocytes proliferate and increase expression of inflammatory factors such as glial fibrillary acidic protein (GFAP) which can later lead to astroglia scar formation along the infarct border (Pekny and Nilsson 2005, Nowicka et al. 2008, Sanchez-Bezanilla et al. 2021). These interactions become even more complex with the understanding that peri-infarct regions in relationship to the core lesion demonstrate distinct mechanisms in a spatial manner. The core region demonstrates progressive cavitation, while the peri-infarct tissue becomes increasingly plastic. Thus, understanding the complexities of these interacting systems as they develop after ischemia in a spatial manner is key to further elucidating the underlying mechanisms for future preventions and treatments.

Gene and protein profile expressions have often been analyzed after ischemic stroke to elucidate these mechanisms during various timepoints (Surles-Zeigler et al. 2018, Xu et al. 2005, Ford et al. 2006, Xu et al. 2004, Rodriguez-Mercado et al. 2012, Simmons et al. 2016, Wang et al. 2015, Xu et al. 2006, Noll et al. 2019), but have been limited by technological challenges. Technology limitations have included the requirement that tissue to be analyzed in sections, such as the entire ipsilateral and contralateral hemispheres. Some studies have carefully extracted cortices and subcortices from each hemisphere with more specific profile success. However, the spatial profile specific to the mechanisms occurring in these distinct regions post-ischemia: the ischemic core, peri-infarct, and peri-infarct normal tissue (PiNT) have not yet been clearly elucidated in such detail. In this study, we examined the spatial regulation of protein profiles initiated by ischemia. By utilizing immunohistochemistry and Nanostring Digital Spatial Profiling (DSP) we analyzed panels of proteins in regions of interest including the ischemic core border, peri-infarct, and PiNT at 3- days post-ischemia compared to regions in the contralateral cortex. NanoString’s GeoMx DSP technology generates profiling data for validated protein analytes using high-plex spatial profiling to rapidly and quantitatively assess the biological implications of the heterogeneity within the tissue samples. We demonstrated that the ipsilateral hemisphere particularly initiated distinct proteomic regulatory profiles in a spatial manner that can be co-localized consistently with the cellular markers, fluoro Jade B (FJB), GFAP, and Iba-1. Additionally, the core border region presented a unique proteomic profile that represents neuronal death, apoptosis, immunoreactivity, and early degeneration. Most notably, the core border resulted in a decrease of the neuronal proteins Map2 and NeuN, an increase in the autophagic proteins BAG3 and CTSD, an increase in the microglial and peripheral immune invasion proteins Iba1, CD45, CD11b, and CD39, and an increase in the neurodegenerative proteins BACE1, APP, αβ 1-42, ApoE, and hyperphosphorylated tau protein S-199. The peri-infarct region demonstrated increased astrocytic immunoreactivity, apoptotic, and neurodegenerative proteomic profile, with an increase in the autophagic protein BAG3, GFAP, and hyperphosphorylated tau protein S-199. The PiNT region displayed minimal changes compared to the contralateral cortex with only an increase in GFAP. These findings may aid in identifying a novel therapeutic strategy for the treatment of stroke.

## Materials and Methods

### Animals

All animals used in these studies were treated humanely and with regard for alleviation of suffering and pain, and all protocols involving animals were approved by the IACUC of University of California-Riverside prior to the initiation of experimentation. Male and female C57BL6 mice (8–10 weeks old) were purchased from Jackson Laboratories (Bar Harbor, Maine) and housed with a 12-hour daily light/dark cycle. Food and water were provided ad libitum. All surgical procedures were performed by sterile/aseptic techniques in accordance with institutional guidelines.

### Photothrombotic Middle Cerebral Occlusion

Animals were subjected to left photothrombotic MCA occlusion (MCAO). Mice were anesthetized with 2% isoflurane and circulating air (N_2_O:O_2_ at approximately 2:1) and maintained anesthetized during the procedure via a modified gas tubing nose connection on a stereotaxic instrument. Eye lubricant was applied to protect the eyes and body temperature was maintained via a heating pad placed underneath the mice during surgery at 37°C. MCAO was performed in an adapted accordance to Zhong et al. (Zhong et al. 2010). Rose Bengal (10 mg/mL; Cat#330000, Sigma, Burlington, MA) was injected intraperitoneally (i.p.) at 10 mL/g and allowed to incubate for 8 minutes. The animal was stabilized in the stereotaxic instrument and the scalp hair was removed. The scalp was disinfected with iodine and ethanol, then followed by a midline skin incision to expose the skull above the left sensorimotor cortex. A 2-mm diameter focal green laser (520nm; Cat# LP520-MF100 ThorLabs Inc, Newton, NJ) was directed at 2-mm lateral left and 0.6-mm posterior of bregma. Laser irradiation occurred for 20 minutes at 10 mW. After irradiation, the midline incision was sealed with Vetbond glue followed by triple antibiotic ointment. Mice were then placed into a 37°C incubation chamber for 20-30 minutes to recover. Sham control animals included two groups: 1) Rose Bengal injection without laser irradiation and 2) laser irradiation without Rose Bengal injection. Mice were sacrificed at 3 days post-ischemia (dpi).

### Histology and Immunohistochemistry

After MCAO, mice were deeply anesthetized with 2% isoflurane and perfused transcardially with saline followed by cold 4% paraformaldehyde (PFA) solution. Brains were quickly removed and maintained in 4% PFA for 24 hours. Brains in preparation for histological and immunohistochemical analysis were quickly removed after transcardial perfusion and maintained in 4% PFA for 24 hours before being cryoprotected in 30% sucrose. The brains were then flash frozen and stored at – 80°C until sectioning. Coronal sections of 12–15 μm thickness were cryosectioned and mounted on slides which were then stored at −80°C until further processed.

Cresyl violet (Cat#C5042, Millipore, Billerica, MA) stain was first reconstituted from powder with distilled water, allowed to stir overnight, and then 0.3% glacial acetic acid was added and mixed thoroughly. Sections were stained with cresyl violet on slides beginning with rehydrating steps of 15 minutes incubation with 95% ethanol, 1 minute with 70% ethanol, 1 minute with 50% ethanol, 2 minutes with distilled water, and 1 minute with distilled water. Sections were then stained with cresyl violet warmed to 37.5°C for 3 minutes followed by distilled water for 1 minute. Sections were then dehydrated with 1 minute of 50% ethanol, 2 minutes of 70% ethanol with 1% glacial acetic acid, 2 minutes of 95% ethanol, and 1 minute of 100% ethanol. After allowing the ethanol to dry off the sections, they were incubated with warmed cresyl violet again for 2 minutes followed by distilled water for 2 minutes. Sections were cleared with a 5-minute wash of Histoclear and mounted and cover slipped with DPX (Cat#06522, Millipore, Billerica, MA).

Fluoro Jade B (FJB; Cat#AG310, Millipore, Billerica, MA) labeling was performed in an adapted accordance to Noll et al. 2019 to ensure infarct presence and record infarct size and location (Noll et al. 2019). Sections were post-fixed with 10% formalin for 10 minutes and then washed twice with PBS for 5 minutes. Sections were then directly incubated in 0.06% potassium permanganate (KMnO_4_) for 3 minutes followed by distilled water for 2 minutes. Sections were then incubated in a freshly prepared solution of 0.0004% FJB with DAPI for 20 minutes, rinsed in distilled water 3 times for 2 minutes, and then dried at 50°C.

For immunohistochemical studies, sections were dried at room temperature for 30 minutes. After rinsing with 0.01M PBS, sections were blocked in PBS containing 5% normal donkey serum and 0.1% Triton X-100 for 1-2 hours at room temperature, rinsed with PBS/0.2% Tween-20 (PBST) and then incubated overnight at 4°C with primary antibodies of polyclonal rabbit anti-Iba-1 (1:1000, Cat#019-19741, Wako, Osaka, Japan) and Cy3-conjugated monoclonal mouse anti-GFAP (1:400, Cat#C9205, Sigma, Burlington, MA). Sections were washed 3 times with PBST, incubated with respective AlexaFluor594-conjugated donkey anti-rabbit IgG antibody (1:400, Cat#711-585-152, Jackson ImmunoResearch Laboratory, West Grove, PA) for 1 hour at room temperature, then rinsed 4 times with PBST before mounting with DAPI-Fluoromount-G (Cat#OB010020, Fisher Scientific, Pittsburgh, PA).

### Immunohistochemical Quantification

A Leica MZ FL III stereo microscope with a tethered dSLR camera was used to capture all digital images of cresyl violet stained sections. A Leica DM5500 B Automated Upright fluorescence microscope was used to capture all digital images of FJB stained sections at x5 magnification. A Nikon TS2-S-SM inverted fluorescence microscope equipped with a CCD camera was used to capture all digital images of sections at ×10 magnification. Cresyl violet images were captured to demonstrate photothrombotic damage at each investigated stereotaxic location (bregma +2, +1, and 0) and each investigated region of interest (ROI) with highlighted focus on the core border region (Figure 1).

**Figure 1.**
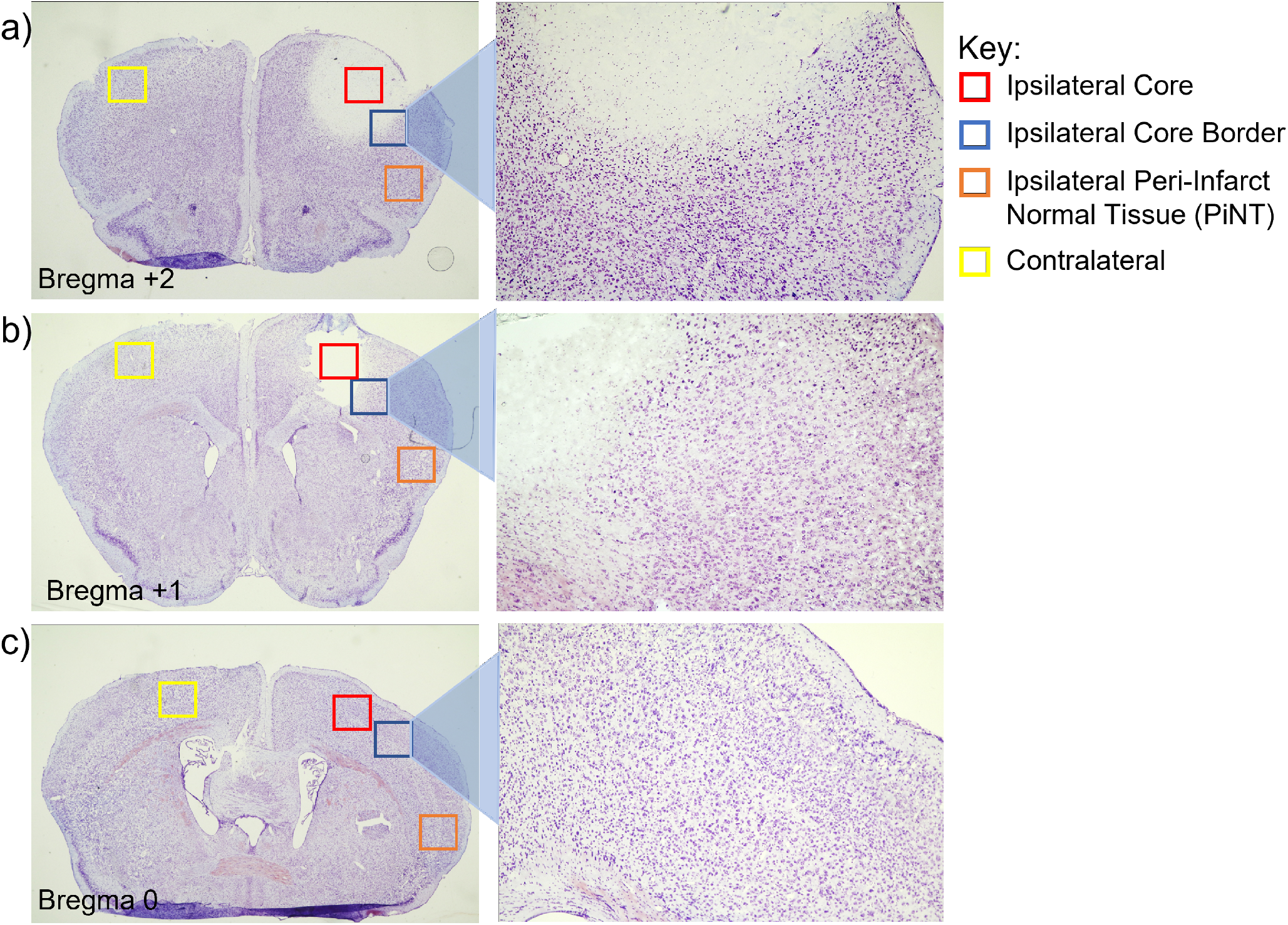
Representative Stereotaxic Locations and Regions of Interest at 3-days post-Ischemia. Coronal sections of mice with induced photothrombotic MCAO were stained with cresyl violet at 3 days after MCAO and examined qualitatively at bregma +2 (a) bregma +1 (b) and bregma 0 (c) with four ROIs: Ipsilateral core (red), Ipsilateral core border (blue), Ipsilateral Peri-Infarct Normal Tissue (PiNT; orange), and Contralateral (yellow) with higher magnification of the core border region. Infarct at 3-days post-ischemia demonstrated larger core damage at bregma +2 with some sustained damage at bregma +1, but no indicated damage at bregma 0. Core border region demonstrates a small increase in cell numbers at both bregma +2 and +1

Immunohistology images of FJB+ cells were captured at bregma +2 and +1 at 5x magnification to capture the entire image area of FJB+ cells. Images were taken at the ipsilateral core and corresponding contralateral cortical region of three separate tissue sections for each mouse. FJB+ cells were counted semi-automatically with ImageJ software (Media Cybernetics, Inc., Bethesda, MD) after threshold at size 10-infinity (pixel units) and circularity 0.4-1.00. FJB+ cell counts were averaged for each stereotaxic location.

Immunohistology images of Iba-1, and GFAP were captured at approximately bregma +2, +1, and 0 at four different regions of interest: the core, core border, PiNT and contralateral cortex in three separate corresponding tissue sections for each mouse. Mean gray values were calculated with ImageJ software. All picture properties were obtained and ensured for consistency. Pictures were converted to 32-bit gray before analysis, and pictures were not altered in any other way. Area fraction was utilized as a control where all picture values must have 100% area fraction. Mean gray value regions of interest technical replicates were averaged and were calculated as fold changes against the contralateral side. Iba-1 maxima count was analyzed with prominence=15, in areas of high tissue damage (based on cresyl violet staining) where background signal may be higher, prominence was increased to 18. Maxima count ROI technical replicates were averaged and calculated as fold changes against the contralateral side. Sham and MCAO groups fold-change mean gray values were compared using Student’s two sample t-tests assuming unequal variances.

### Formalin-Fixed Paraffin Embedded Tissue Preparation

Brains in preparation for Nanostring DSP were continually dehydrated in preparation for paraffin embedding: 24 hours of 4% PFA was followed by 24 hours each in 40% ethanol, 70% ethanol, and a second change of 70% ethanol. Brains were then placed whole into cassettes and processed in a Tissue-Tek Processor in a 12-hour cycle (Table 1) before embedding coronally in paraffin.

**Table 1.**
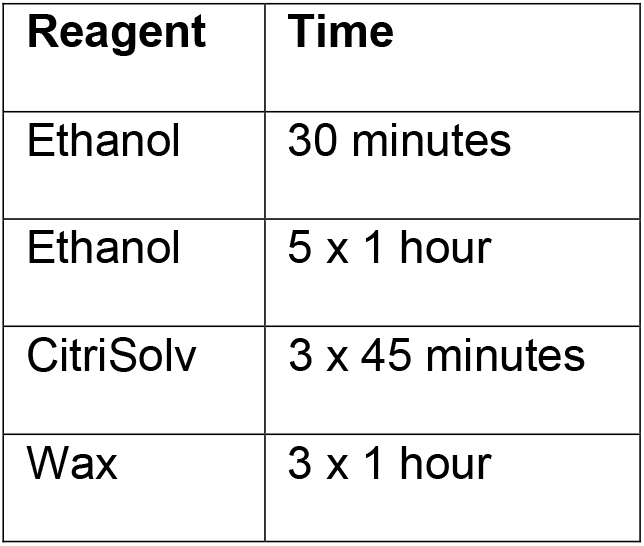
Tissue-Tek Processing Protocol

Formalin-fixed, paraffin-embedded (FFPE) samples were sectioned to approximately bregma +2 and 5-μm thickness sections were mounted onto slides and allowed to dry at room temperature overnight. FJB labeling was performed as described above to ensure infarct presence and record infarct size and location. Samples confirmed with FJB+ cells indicating successful MCAO were sent to Nanostring (Seattle, WA) for slide mounting and GeoMx DSP profiling.

### Nanostring Digital Spatial Profiling Analysis

To visualize whole tissue, mounted slides were stained with oligo-conjugated antibodies for MAP2, Iba-1 and GFAP, and with Syto13 (nuclei) in the GeoMx DSP instrument. Circular geometric patterns (200 μm diameter) were used to identify six ROIs on the scanned tissues of each sample: core border, peri-infarct, and PiNT regions on ipsilateral and contralateral hemispheres. The GeoMx Neural Cell Profiling protein panel (73 proteins) was utilized for this analysis. UV-cleavable oligo-conjugated antibodies according to the panel were dispensed onto each ROI, UV-cleaved off, aspirated into a plate, hybridized and counted by the GeoMx DSP instrument. All resulting spatial, quantified analysis was performed in the Nanostring GeoMx DSP analysis software. Background correction was determined by protein target correlation plot and high correlation was seen between Rb IgG, Rb IgGa, and Rb IgGb. All three IgG proteins were used for panel background correction via signal-to-background ratio. Spatial ROIs were compared using a linear mixed model (LMM) with Bonferroni-Hochberg (BH) correction and random effect for Scan ID and ROI ID. Fold changes were identified by comparing the ROI/contralateral ROI with a significance of p value <0.05.

## Results

### FluoroJadeB Identifies Ischemic Infarct 3 Days Post-MCAO

We examined the consistency and ability to define the photothrombotic infarct at three days post-injury with the neurodegeneration stain, FJB. Brain tissues from mice with induced MCAO were sacrificed and examined spatially at three days after ischemia at stereotaxic locations bregma +2 and +1. Detectable FJB+ cells were found consistently throughout the infarct core at bregma +2, but not at bregma +1 (Figure 2). MCAO brain tissue exhibited a mean of 547.3 ± 114 FJB+ cells at bregma +2 with only a mean of 11.6 ± 19 at bregma +1. This demonstrated that neurodegenerative damage occurs and is detectable by FJB staining three days after MCAO with primary neuronal damage located at bregma +2.

**Figure 2.**
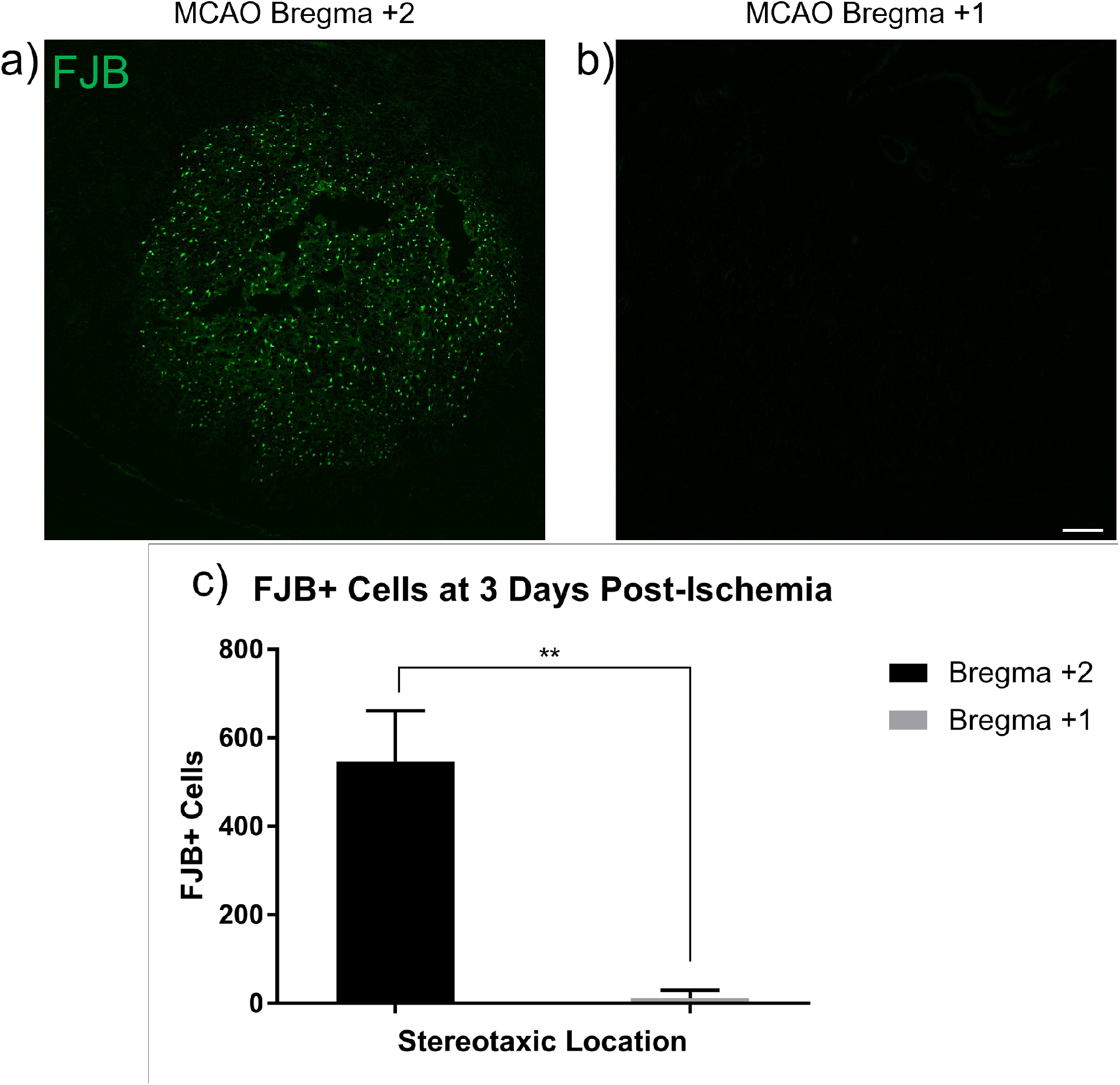
Neuronal Damage 3 Days Post-Ischemia. Brains from mice with induced MCAO were examined 3 days after ischemia. FJB+ cells were counted at bregma +2 and +1 at 5x magnification for whole infarct comparison. Extensive FJB+ staining was found at bregma +2 (a) but were not detected at bregma +1 (b). A mean value of FJB+ cells from three brain sections from each location was obtained for each individual mouse brain (c; p<0.01, n=4). Data are expressed as mean ± SD. Scale bar = 100 µm.

### GFAP Increases in Core Border 3 Days Post-Ischemia

To characterize the early inflammatory processes that occur after MCAO, astrocytic immunoreactivity was assessed in a spatial manner with the immunohistology marker, GFAP. Brain tissues from mice with induced MCAO and sham controls (SHAM) were sacrificed and examined at three days after ischemia at bregma +2, +1, and 0 within the ipsilateral core, core border, and PiNT for GFAP expression. High GFAP expression is not typically observed in the naïve or SHAM mouse cortex but is upregulated within proliferating and activated astrocytes following neuronal injury (Davies et al. 1998, Savchenko et al. 2000, Nowicka et al. 2008). Therefore, GFAP expression was analyzed as mean gray value fold change against the contralateral side to detect regional changes in GFAP expression. GFAP regional fold change expression in MCAO animals was compared to SHAM animals (Figure 3). GFAP expression significantly increased within the ipsilateral core border at both bregma +2 and +1 with a fold change of 1.76 ± 0.3 and 1.66 ± 0.29, respectively. No significant changes in GFAP mean gray value fold changes between MCAO and SHAM were seen in ipsilateral core or PiNT regions at any stereotaxic location. Additionally, GFAP mean gray fold change expression was non-distinct from SHAM at bregma 0 for all three regions. GFAP expression suggests that there is an increase in astrocytic inflammatory activity along the core border in a spatial manner at three days post-ischemia.

**Figure 3.**
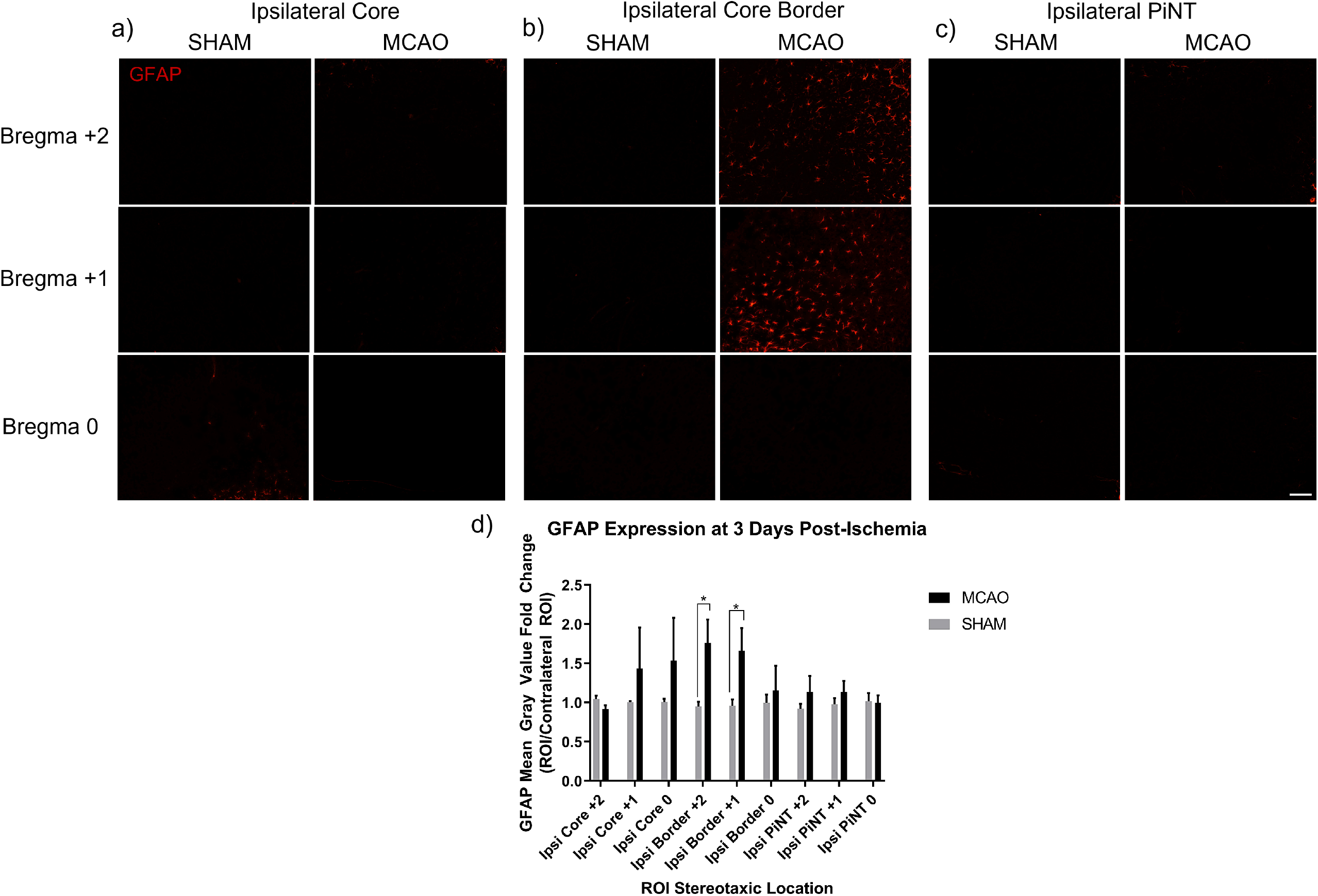
GFAP Expression Increases in Core Border Spatially 3 Days Post-Ischemia. Brains from mice with induced photothrombotic MCAO were examined spatially 3 days after MCAO for GFAP expression. Mean gray value fold change of GFAP expression compared to the contralateral hemisphere in the ipsilateral core (a) ipsilateral core border (b) ipsilateral PiNT (c). Mean gray value fold change increased significantly in the core border at bregma +2 and bregma +1 but was non-significant in bregma 0 (d). Mean gray value fold change of GFAP was non-significant in ipsilateral core or PiNT in any stereotaxic location. Data are expressed as mean ± SD; p<0.05, MCAO (n=4) and SHAM (n=2). Scale bar = 100 µm.

### Microglia Increase Reactivity Within Ischemic Region

To further characterize the inflammatory processes that occur after MCAO, microglial activity was assessed in a spatial manner with the immunohistology marker, Iba-1. Brain tissues from mice with induced MCAO and SHAM surgery were sacrificed and examined spatially at three days after ischemia at bregma +2, +1, and 0 within the ipsilateral core, core border, and PiNT for Iba-1 expression. In the SHAM mouse cortex, Iba-1 expressing microglia exhibit a quiescent, ramified morphology (Figure 4) (Nowicka et al. 2008). After ischemia, resident microglia can increase Iba-1 reactivity, change morphology, as well as infiltrating monocytes will express Iba-1 (Ito et al. 2001). Iba-1 expression was analyzed as mean gray value fold change and gray maxima count fold change against the contralateral side to detect regional changes in Iba-1 expression. Iba-1 regional fold changes and gray maxima count fold changes in MCAO animals were compared to SHAM animals (Figure 4). Iba-1 mean gray value fold change significantly decreased within the core region at stereotaxic location bregma +2 with a fold change of 0.74 ± 0.09, a -1.3 fold change compared to SHAM (Figure 4d). Iba-1 gray maxima count also significantly decreased within the core region at bregma +2 with a fold change of 0.28 ± 0.2, a -3.6 fold change to SHAM (Figure 4e). Iba-1 gray maxima count fold change also exhibited a significant increase compared to SHAM within the core region at bregma 0 with a fold change of 1.35 ± 0.2. Immunohistology images (Figure 4a) demonstrate a distinct morphological change in Iba-1 expressing cells within the bregma +2 core region. There was also a disappearance of Iba-1 expressing cells from the center of the core region, likely reflecting within the decrease of gray values, specifically maxima count. Additionally, round Iba-1 expressing cells appear along the core border region with beginning invasion into the core, suggesting activation of residential microglia and potentially early invasion of peripheral monocytes.

**Figure 4.**
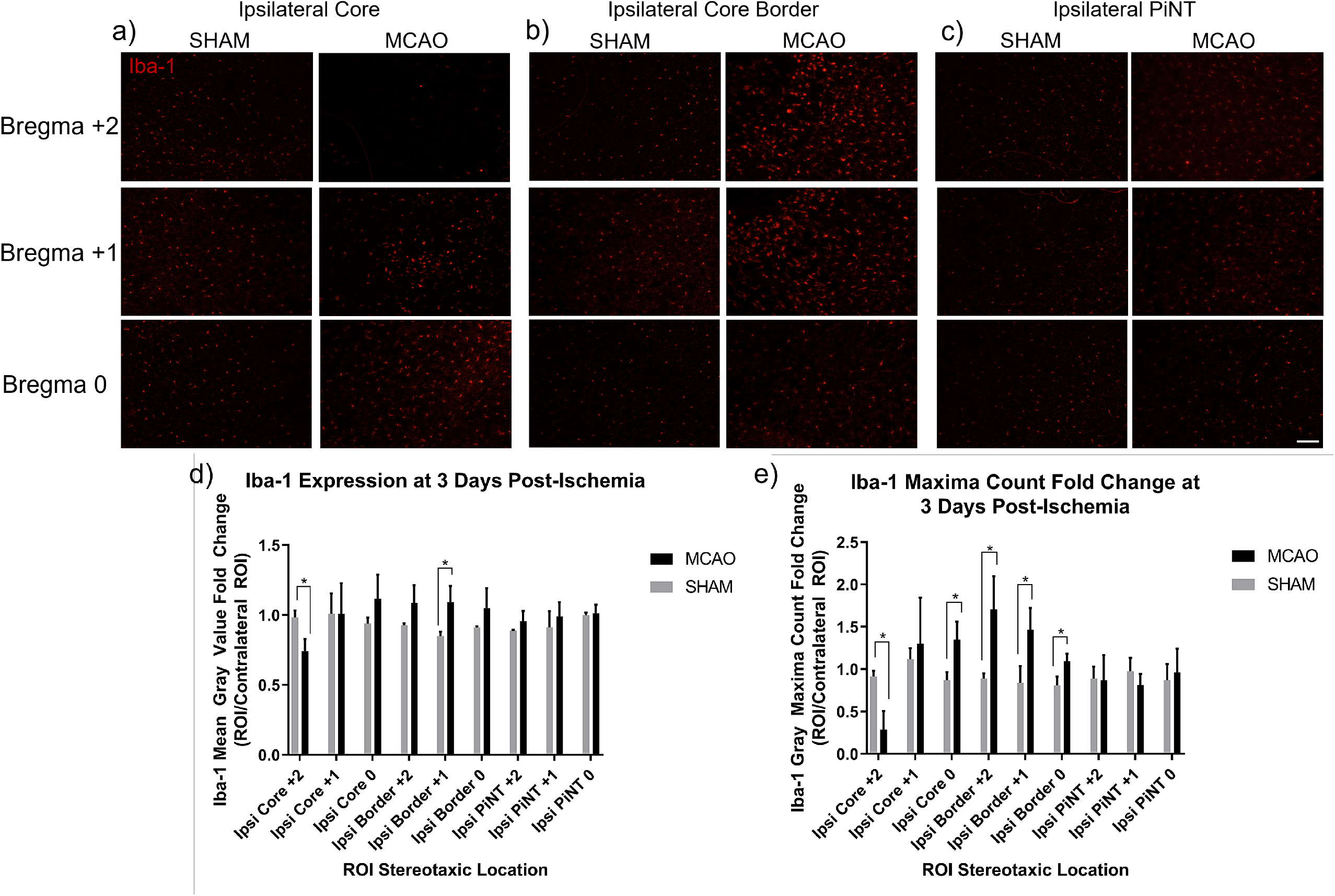
Iba-1 Expression Demonstrates Microglia Reactivity Spatially at 3 Days Post-Ischemia. Brains from mice with induced MCAO were examined spatially 3 days after ischemia for Iba-1 expression. Mean gray value fold change and gray maxima count fold change of Iba-1 expression compared to the contralateral hemisphere in the ipsilateral core (a) ipsilateral core border (b) and ipsilateral PiNT (c). Mean gray value (d) fold change decreased in core region at bregma +2 and increased in the core border at bregma +1. Mean gray value fold change of Iba-1 was non-significant in PiNT or in other regions of interest at any other stereotaxic location. Gray maxima count fold change of Iba-1 decreased in ipsilateral core region at bregma +2 but increased in the core region at bregma 0 and all stereotaxic ipsilateral core border regions (e). Gray maxima count fold change of Iba-1 was non-significantly different in PiNT at all stereotaxic locations. Data are expressed as mean ± SD; p<0.05, MCAO (n=4) and SHAM (n=2). Scale bar = 100 µm.

Iba-1 mean gray value expression minimally increased within the core border at bregma +1 with a fold change of 1.09 ± 0.12 compared to SHAM (Figure 4d). Iba-1 gray maxima count fold change demonstrates significantly increased Iba-1 counts at all core border stereotaxic bregma locations (Figure 4e). Iba-1 gray maxima count fold change within the core border at bregma +2, bregma +1, and bregma 0 were 1.71 ± 0.4, 1.47 ± 0.2, and 1.1 ± 0.1 respectively. The increase in gray maxima count Iba-1+ round cells within each bregma core border location further suggests activation of residential microglia and early invasion of peripheral monocytes into the core in a spatial manner at three days post-ischemia. No significant changes in Iba-1 mean gray value fold changes between MCAO and SHAM were seen in ipsilateral core or core border regions at any other stereotaxic location. Additionally, Iba-1 mean gray fold change expression was non-distinct from SHAM at bregma 0. No significant changes in Iba-1 gray maxima count fold change were seen in the PiNT at any stereotaxic location. Inflammatory activation within the core border of GFAP- and Iba-1- expressing astrocytes and microglia demonstrates the importance of this region early after ischemia. Therefore, we have determined that consistent identification of the core border can be histologically defined with combination of these three glia cell stains: FJB, GFAP, and Iba-1 at three days post-ischemia (Figure 5).

**Figure 5.**
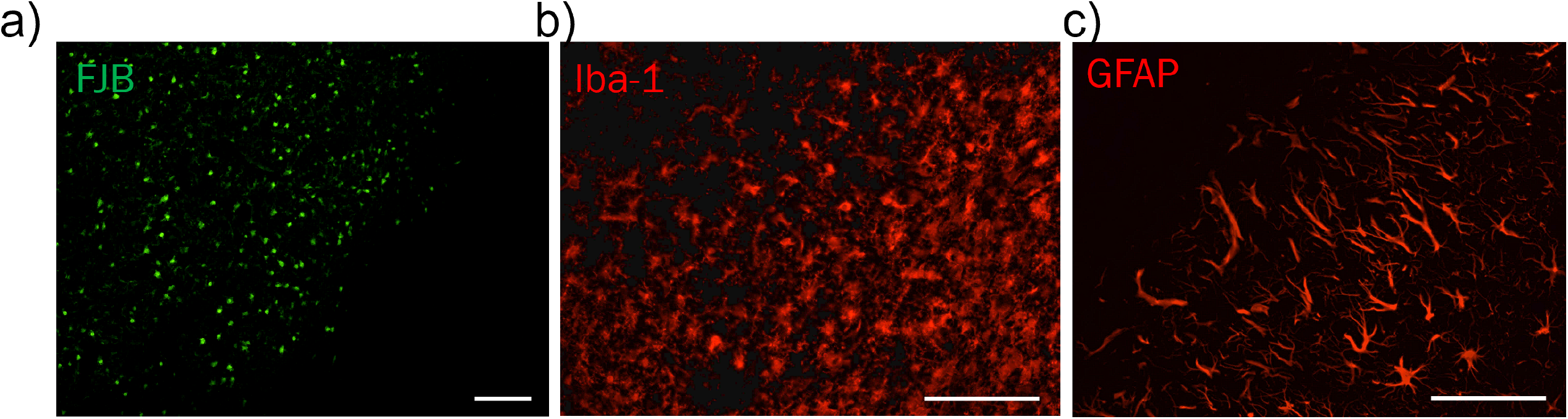
Histological Markers Define Core Border at 3 Days Post-Ischemia. Immunohistology images of FJB (a; green), Iba-1 (b; red), and GFAP (c; red) compared at the core border at bregma +2 can define a consistent, clear core border at 3 days post-ischemia. Scale bar = 100 µm.

### Nanostring Digital Spatial Profiling Defines Distinct MCAO Protein Profiles

In order to elucidate a detailed proteomic analysis and profile after ischemia within these specific ROI’s, we utilized Nanostring’s DSP technology. Brain tissues from mice with induced MCAO were sacrificed and examined spatially at three days after ischemia in the ipsilateral core border, peri-infarct, and PiNT tissues compared to the contralateral side with the Nanostring DSP Neural Cell Profiling protein panel (70 proteins; excluding IgG negative targets). Sections were immunostained with four chosen identifying markers for ROI selection (Figure 6). Markers included: MAP2 for neuronal damage, GFAP for astrocytic reactivity, Iba-1 for microglia, and Syto13 for nuclei. The full neural cell profiling protein panel was run on each ROI resulting in quantified protein read-outs for each ROI.

**Figure 6.**
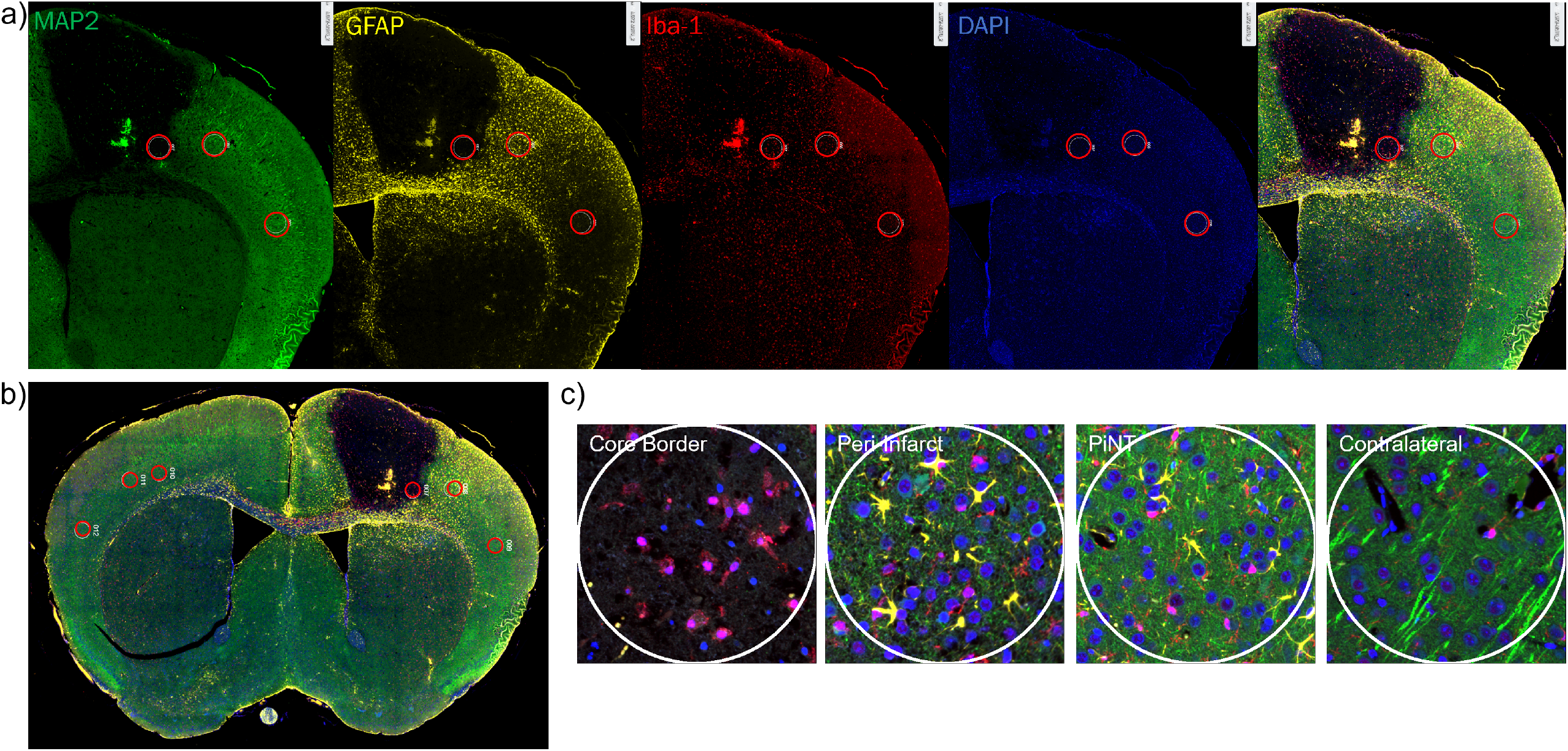
Representative Immunohistochemical Figures for DSP Analysis and ROI Selection. Coronal sections from mice with induced photothrombotic MCAO were immunohistologically stained for identifying markers MAP2 (green), GFAP (yellow), Iba-1 (red), and Syto13 for nuclei (blue) (a) and overlayed for ROI selection (b). Overlay of these markers clearly identified the extent of the infarct region and astroglia immunoreactivity within the core border. ROI’s included ipsilateral core border, ipsilateral peri-infarct, ipsilateral PiNT, and contralateral cortex, each region is outlined in a red circle (c). Clear difference in the identifying IHC markers can be seen within each ROI.

To determine the comparative significance of DSP regional specific comparisons to whole tissue analysis, DSP proteomic profiles were first compared as a combined ipsilateral hemispheric group to contralateral hemispheres proteomic data. All ipsilateral ROIs (core border, peri-infarct, and PiNT ROIs) were combined as one ipsilateral hemispheric group and compared against the contralateral side for significantly changed proteins via an LMM. Then, the individual core ROI was compared against the contralateral side for significantly changed proteins via an LMM (Figure 7). The whole ipsilateral hemispheric group resulted in only 4 significantly upregulated proteins. However, when the ipsilateral core border ROI was isolated and compared to the contralateral tissues, 27 proteins demonstrated significant changes, with 19 upregulated and 8 downregulated.

**Figure 7.**
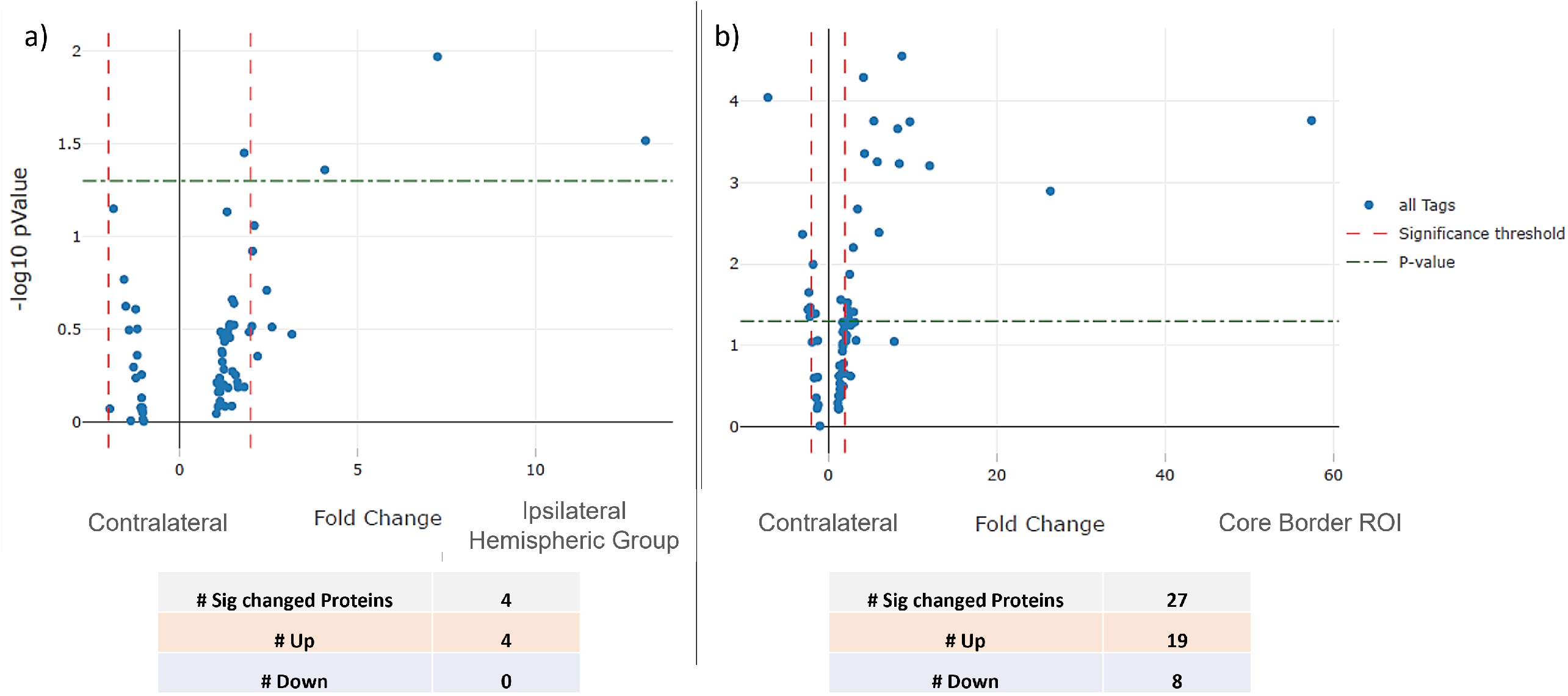
Ischemia Presents Region Specific Proteomic Profiles. Volcano plots representing the significant protein fold changes of the Ipsilateral Hemispheric group (core border, peri-infarct, and PiNT ROIs) (a) and isolated core border ROI (b) compared against the contralateral ROI. Data is plotted on a -log10 pValue scale; fold change on the x-axis; all tags (blue plotted dots) represent individual panel proteins; p-value (0.05; green dotted x parallel line) (n=3).

To determine general similarities and consistencies between the proteomic profiles of ROIs between biological samples, a hierarchical cluster was created (Figure 8). Overall, all core border ROIs clustered together, demonstrating the most similar profiles across samples within the core border. Peri-infarct ROIs clustered closest to the core border ROIs, demonstrating similarity between samples within the peri-infarct region and closest proteomic profile next to the core border ROIs. The PiNT and contralateral ROIs clustered generally together outside of the core border and peri-infarct regions, demonstrating most similarity to each other between samples and furthest similarity to the core border and peri-infarct regions. Overall, this hierarchal cluster suggests that each ROI group demonstrates a unique, consistent proteomic profile after ischemia.

**Figure 8.**
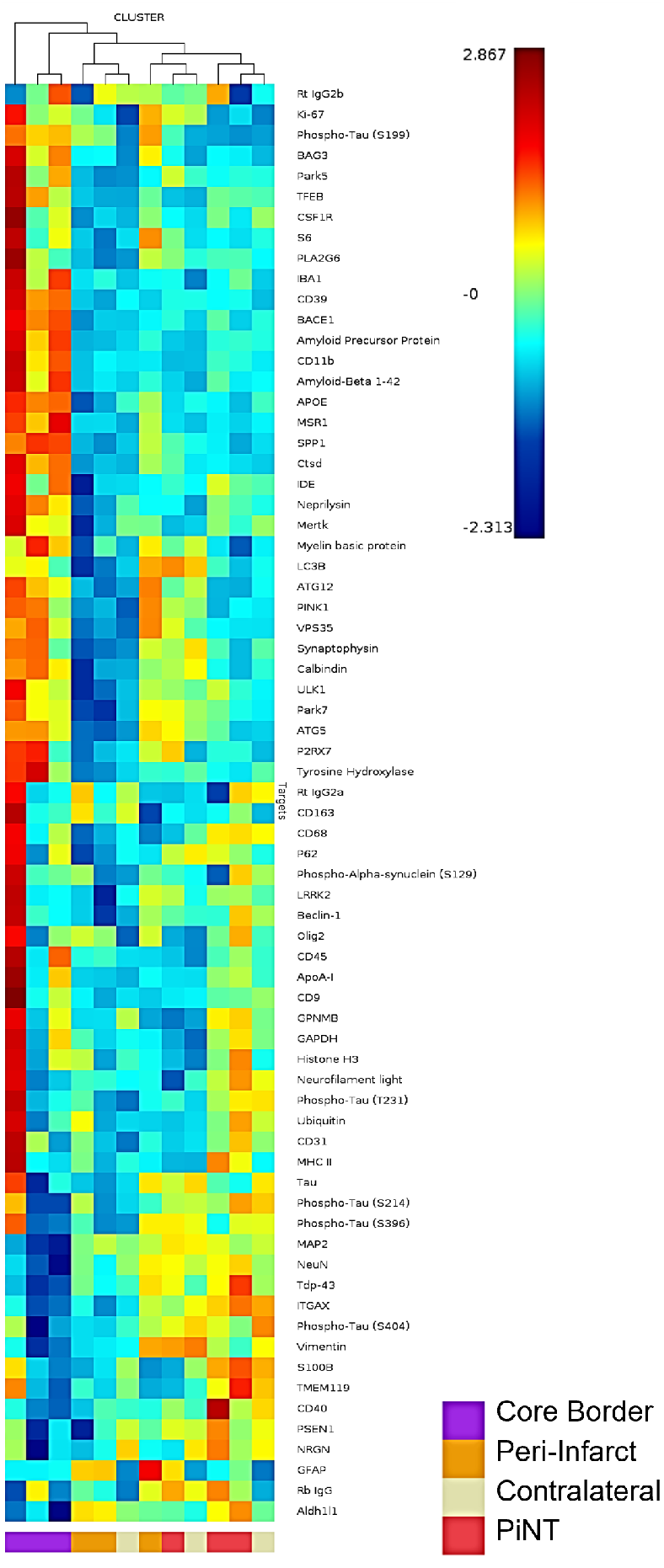
Hierarchical Cluster Heat Map of ROIs. Protein expression from each individual ROI representing the ipsilateral core border (purple), ipsilateral peri-infarct (orange), ipsilateral PiNT (red), and contralateral (white) is plotted on a heat map where red= higher representative expression and blue= lower representative expression. ROIs are clustered according to proteomic expression similarity where most similar are clustered together and tree relatives are mapped above.

Each ROI was further characterized compared to the contralateral ROI. Significantly differentially regulated proteins according to fold change were documented for each ROI (Table 2). The ipsilateral core border ROI demonstrated 27 differentially regulated proteins with 19 upregulated and 8 downregulated. Of these 27 differentially regulated proteins, there was most notably a unique profile reflected in neuronal death, apoptotic, immunoreactive, and early degeneration proteins. Neuronal proteins, MAP2 and NeuN resulted in downregulation of -7.19 and -3.07 respectively in the core border. An autophagy protein, BAG3, resulted in a 12.04-fold upregulation and a lysosomal autophagy protein, CTSD, was also upregulated by 8.44-fold. The astrocytic protein, Aldh1/1, was downregulated by -2.30-fold and a glial cytoskeletal intermediate filament protein, vimentin, was downregulated by -2.15-fold. Many proteins related to microglial immunoreactivity and immune peripheral invasion were significantly upregulated within the core border region. These included a 2.25-fold upregulation in CD45, a 4.31-fold upregulation in Iba-1, an 8.24-fold upregulation in CD11b, an 8.74-fold upregulation in CD39, and a 9.69-fold upregulation in MSR1. Notably, there was also a marked 26.39-fold upregulation in SPP1, a protein specifically associated with disease-related microglia.

**Table 2.** Regulatory Proteomic ROI Profiles. Proteins were compared against the contralateral side for significant fold changes via an LMM for each ROI including: Ipsilateral Hemispheric Group, ipsilateral core border ROI, ipsilateral peri-infarct ROI, and ipsilateral PiNT ROI. Upregulated proteins expressed in yellow-green, downregulated proteins expressed in orange-red (p<0.05, n=3).

|  |  | Ipsilateral Hemispheric Group | Ipsilateral Core Border ROI | Ipsilateral Peri-Infarct ROI | Ipsilateral PiNT ROI |
| --- | --- | --- | --- | --- | --- |
|  | *Differentially Regulated Proteins* | 4 | 27 | 3 | 1 |
|  | *Upregulated* | 4 | 19 | 3 | 1 |
|  | *Downregulated* | 0 | 8 | 0 | 0 |
| Protein Name | Protein Acronym | Fold Change | Fold Change | Fold Change | Fold Change |
| Microtubule associated protein 2 | MAP2 |  | -7.19 |  |  |
| Neuronal nuclei | NeuN |  | -3.07 |  |  |
| BAG family molecular chaperone regulator 3 | BAG3 | 4.08 | 12.04 | 2.96 |  |
| Cathepsin D | CTSD |  | 8.44 |  |  |
| Transcription factor EB | TFEB |  | 2.96 |  |  |
| Vacuolar protein sorting ortholog 35 | VPS35 |  | 2.36 |  |  |
| Aldehyde dehydrogenase 1 family member A1 | Aldh1l1 |  | -2.30 |  |  |
| Vimentin | VIM |  | -2.15 |  |  |
| Glial fibrillary acidic protein | GFAP | 7.25 |  | 23.69 | 6.09 |
| Cluster of differentiation 45 | CD45 |  | 2.25 |  |  |
| Ionized calcium binding adaptor molecule 1 | IBA1 |  | 4.31 |  |  |
| Cluster of differentiation molecule 11B | CD11b |  | 8.24 |  |  |
| Cluster of differentiation 39 | CD39 |  | 8.74 |  |  |
| Macrophage scavenger receptor 1 | MSR1 |  | 9.69 |  |  |
| Secreted phosphoprotein 1 | SPP1 |  | 26.39 |  |  |
| P2X purinoceptor 7 | P2RX7 | 1.82 | 3.01 |  |  |
| Complement component 3 receptor 4 subunit | ITGAX |  | -1.53 |  |  |
| Beta-secretase 1 | BACE1 |  | 4.18 |  |  |
| Amyloid Precursor Protein | APP |  | 5.41 |  |  |
| Amyloid-Beta 1-42 | αβ 1-42 |  | 5.82 |  |  |
| Phospho-Tau (S199) | tau-S199 | 13.10 | 57.42 | 21.41 |  |
| Phospho-Tau (S404) | tau-S404 |  | -2.19 |  |  |
| Apolipoprotein E | ApoE |  | 6.02 |  |  |
| Insulin-degrading enzyme | IDE |  | 1.48 |  |  |
| Neprilysin | NEP |  | 3.48 |  |  |
| TAR DNA-binding protein 43 | Tdp-43 |  | -1.81 |  |  |
| Neurogranin | NRGN |  | -2.42 |  |  |
| Parkinson Disease 5 | Park5 |  | 2.54 |  |  |

Within the core border region, BACE1 was upregulated by 4.18-fold, APP was upregulated by 5.41-fold, and amyloid β (αβ) 1-42 was upregulated by 5.82-fold. This suggests that three days after ischemia, the BACE1 cleaving protein was upregulated within the core and perhaps leads to increased cleaved products of APP and the αβ 1-42 form. Notably, there was a significant upregulation of the tau-S199 amyloid by 57.42-fold, which can lead to neuronal destabilization and death. There was also a significant upregulation of ApoE by 6.02-fold, which could later lead to be neuroprotective or neurodegenerative depending on whether it is expressing the neuroprotective isoform 3 or neurodegenerative isoform 4 (Koistinaho and Koistinaho 2005). The ipsilateral peri-infarct ROI demonstrated significant upregulation in 3 proteins: GFAP, was upregulated by 23.69-fold, tau-S199 was upregulated by 21.41-fold, and BAG3 was upregulated by 2.96-fold. The ipsilateral PiNT ROI only demonstrated significant upregulation in one protein, GFAP, by 6.09-fold.

## Discussion

Our data presented emphasizes the importance of accounting for spatial differences when analyzing post-ischemic proteomic data. Our proteomic analysis presents unique profiles spatially and regionally in relation to the infarct core. Previous technologies using whole ipsilateral tissues can provide some insight into mechanisms occurring post-ischemia but can greatly obscure specific mechanisms that occur within each region. Our data demonstrated that each region analyzed with DSP demonstrated a unique proteomic profile with distinctly regulated proteins. Within the ischemic core border, a unique profile relating to neuronal death, apoptosis, immunoreactivity, and early degeneration was identified. When looking further into the relationship between these differentially regulated proteins, a potential pattern emerged with future interest for therapeutic targeting. As expected, the neuronal proteins, MAP2 and NeuN were downregulated, indicative of neuronal death within the core border. This is also reflected in the upregulation of the autophagic protein, BAG3, and the lysosomal autophagy protein, CTSD. CTSD has been shown to increase sharply after ischemia but will decrease as lysosomes rupture (Liu et al. 2019). BAG3 was also upregulated within the peri-infarct region, suggesting that autophagy and related apoptosis may still be ongoing within this region.

Ischemia also resulted in an upregulation of the general immune cell surface marker, CD45, as well as Iba-1. This suggests an increase in residential microglia and/or peripheral monocyte infiltration and activity. However, as residential microglia are recognized to decrease acutely within the core via degeneration, which is reflected within the immunohistochemistry mean gray value and gray maxima count fold changes, we expect this Iba-1 increase to reflect peripheral monocyte invasion (Ito et al. 2001, Matsumoto et al. 2007, Nowicka et al. 2008). This is also suggested by the 8-fold upregulation of CD11b and CD39 and a 3-fold upregulation of the CD39 receptor, P2RX7, which are all surface markers on leukocytes. Leukocytes are noted to begin infiltrating into the ischemic region as early as 12 hours after ischemia and persist, with peak levels at 7 days post-ischemia (Nowicka et al. 2008, Ito et al. 2001, Sanchez-Bezanilla et al. 2021). Microglia reactivity post-ischemia is notoriously recognized to have both beneficial and deleterious effects. Activated microglia acutely can phagocytose cells that have necrotized within the lesion core, preventing the further release of neurotoxic products (Giulian, Vaca and Corpuz 1993b, Giulian et al. 1993a, Matsumoto et al. 2007). However, activated microglia also release many pro-inflammatory cytokines and reactive oxygen species which can contribute to infarct development (Giulian et al. 1993b, Giulian et al. 1993a, Matsumoto et al. 2007). Therefore, truly understanding the interactions of these cytokines and proteins is key to identifying future targets for therapeutic intervention. For example, CD39 and MSR1 both demonstrated an upregulation of approximately 9-fold, and both proteins have been suggested to play a neuroprotective role within the inflammatory process. Transgenic mouse lines overexpressing CD39 resulted in reduced leukocyte infiltration, smaller infarct volumes, and decreased neurological deficit after ischemia, suggesting a neuroprotective role of CD39 during ischemic insult (Baek et al. 2017). CD39 expression on endothelial cells and leukocytes is suggested to reduce inflammatory cell trafficking and platelet reactivity via cell-cell interaction after ischemia (Hyman et al. 2009). Also, it has been shown that higher expression of the MSR1 protein increases damaged-associated molecular patterns (DAMP) clearance after ischemia via MSR1-induced resolution of neuroinflammation, indicating MSR1 as a potential therapeutic target (Zou et al. 2021). Additionally, there was a notable, 57-fold upregulation of the protein SPP1. This protein is suggested to function as a pro-angiogenic trophic factor, but it is also associated with neurodegeneration as it is recorded to be upregulated in senescent microglia in an age-dependent manner (Li et al. 2010, Jiang et al. 2020). Controversially, this protein has also been associated with macrophagic communication to astrocytic migration toward the infarct area that has been suggested to be a neuroprotective repair mechanism of the ischemic neurovascular unit early after ischemia (Choi et al. 2019).

A unique astrocytic proteomic profile is reflective throughout each region. Within the core border, there is downregulation of the astrocytic proteins, Aldh1/1 and vimentin by about 2-fold. Vimentin-expressing astrocytes have been shown to increase acutely after ischemia but given downregulation at this three-day timepoint in combination with Aldh1/1 downregulation, may suggest astrocytic degeneration within the core (Kindy, Bhat and Bhat 1992). However, large upregulation of GFAP within the peri-infarct region by almost 24-fold and confirmed with mean gray value fold change increase at the core border both at bregma +2 and +1 suggests strong reactivity of astrocytes within the border and peri-infarct region three-days after ischemia. GFAP has been shown by multiple studies to be increased specifically within the peri-infarct region beginning as early as 4 hours after ischemia up until 28 days (Kindy et al. 1992, Nowicka et al. 2008, Sanchez-Bezanilla et al. 2021). Reactivity of astrocytes and microglia can lead to secondary tissue damage, progressive cavitation, and scar formation along the peri-infarct border (Fitch et al. 1999). Ultimately, the relative differences between these microglial and astrocytic proteomic profiles within the core border and peri-infarct regions could provide potential targets for prevention of secondary damage and anti-inflammation.

Ischemia is well-recognized to be associated with neurodegeneration and more specifically the relationship between stroke history, tau proteins, and increased Alzheimer’s Disease prevalence (Sun et al. 2006, Zheng et al. 2010). Multiple tau related proteins within the core border region resulted in a pattern that could ultimately indicate a future therapeutic target. BACE1, APP, and αβ 1-42 all demonstrated an approximate 5-fold upregulation within the core border. BACE1 has been shown to be upregulated in response to released reactive oxygen species and hypoxia inducible factor 1α (HIF1α) following ischemia (Sun et al. 2006, Guglielmotto et al. 2009). Increased BACE1 activity results in increased cleavage of APP with increased deposits and aggregation of the neurotoxic form of amyloid β, αβ 1-42 (Sun et al. 2006, Guglielmotto et al. 2009). BACE1 expression was found to be regulated by a positive feedback loop between γ- and β-secretase cleavages on APP (Wang et al. 2006, Li et al. 2009). Additionally, upregulation of APP within neurons, astrocytes, or microglia can lead to neuronal destabilization and vulnerability to stress, suggesting that upregulation of all three proteins within the core border could be indicative of a neurodegenerative profile (Koistinaho and Koistinaho 2005). The tau-S199 form was also found highly upregulated by 57-fold within the core border, and this tau form has been shown to play a role in the development of Alzheimer-like lesions after ischemia. After ischemia, tau proteins become rapidly dephosphorylated acutely and are then slowly rephosphorylated and accumulated in a serine site-specific manner (Zheng et al. 2010). Tau-S199 has been reported to be specifically induced after ischemia and to contribute to ischemic neuronal injury (Zheng et al. 2010). Dysregulation and displacement of tau proteins after ischemia is suggested to contribute to a neurodegenerative profile. Lastly, upregulation of the protein ApoE within the core border region could play a dual role depending on the isoform of ApoE that is being expressed. ApoEε3 is the most common isoform and can exhibit a neuroprotective effect by contributing to communication from microglia to neurons for repair, regeneration and survival as well as suppression of microglia reactivity (Koistinaho and Koistinaho 2005). However, the ApoEε4 isoform may be neurodegenerative by contributing to αβ deposition (Koistinaho and Koistinaho 2005).

## Conclusion

The ability to isolate the ipsilateral core border, ipsilateral peri-infarct, ipsilateral PiNT, and contralateral cortex regions and compare their proteomic profiles in such a detailed manner has allowed us to determine that each region demonstrates a unique differentially regulated proteomic profile. Future ischemia studies could greatly benefit from spatial analysis as we have demonstrated the substantial loss of data that can occur with whole tissue analysis. Overall, our data highlight the importance of identifying ischemic spatial mechanisms to understand the complex, dynamic interactions throughout ischemic progression and repair as well to introduce potential targets for successful ischemic therapeutic interventions.

## Acknowledgements

This study was partly funded by NIH grants to BDF (R01NS091616; R25GM119975; R21NS106949).

## References

Baek, A. E., N. R. Sutton, D. Petrovic-Djergovic, H. Liao, J. J. Ray, J. Park, Y. Kanthi & D. J. Pinsky (2017) Ischemic Cerebroprotection Conferred by Myeloid Lineage-Restricted or Global CD39 Transgene Expression. Circulation, 135, 2389–2402.

Barone, F. C. & G. Z. Feuerstein (1999) Inflammatory mediators and stroke: new opportunities for novel therapeutics. J Cereb Blood Flow Metab, 19, 819–34.

Benjamin (2017) Heart Disease and Stroke Statistics-2017 Update: A Report From the American Heart Association (vol 135, pg e146, 2017). Circulation, 136, E196–E196.

Bosetti, F., J. I. Koenig, C. Ayata, S. A. Back, K. Becker, J. P. Broderick, S. T. Carmichael, S. Cho, M. J. Cipolla, D. Corbett, R. A. Corriveau, S. C. Cramer, A. R. Ferguson, S. P. Finklestein, B. D. Ford, K. L. Furie, T. M. Hemmen, C. Iadecola, L. B. Jakeman, S. Janis, E. C. Jauch, K. C. Johnston, P. M. Kochanek, H. Kohn, E. H. Lo, P. D. Lyden, C. Mallard, L. D. McCullough, L. M. McGavern, J. F. Meschia, C. S. Moy, M. A. Perez-Pinzon, I. Ramadan, S. I. Savitz, L. H. Schwamm, G. K. Steinberg, M. P. Stenzel-Poore, M. Tymianski, S. Warach, L. R. Wechsler, J. H. Zhang & W. Koroshetz (2017) Translational Stroke Research: Vision and Opportunities. Stroke, 48, 2632–2637.

Brouns, R. & P. P. De Deyn (2009) The complexity of neurobiological processes in acute ischemic stroke. Clin Neurol Neurosurg, 111, 483–95.

Choi, I. A., J. H. Yun, J. H. Kim, H. Y. Kim, D. H. Choi & J. Lee (2019) Sequential Transcriptome Changes in the Penumbra after Ischemic Stroke. Int J Mol Sci, 20.

Cramer, S. C., R. Shah, J. Juranek, K. R. Crafton & V. Le (2006) Activity in the peri-infarct rim in relation to recovery from stroke. Stroke, 37, 111–5.

Davies, C. A., S. A. Loddick, R. P. Stroemer, J. Hunt & N. J. Rothwell (1998) An integrated analysis of the progression of cell responses induced by permanent focal middle cerebral artery occlusion in the rat. Exp Neurol, 154, 199–212.

del Zoppo, G., I. Ginis, J. M. Hallenbeck, C. Iadecola, X. Wang & G. Z. Feuerstein (2000) Inflammation and stroke: putative role for cytokines, adhesion molecules and iNOS in brain response to ischemia. Brain Pathol, 10, 95–112.

Detante, O., A. Jaillard, A. Moisan, M. Barbieux, I. M. Favre, K. Garambois, M. Hommel & C. Remy (2014) Biotherapies in stroke. Rev Neurol (Paris), 170, 779–98.

Dirnagl, U., C. Iadecola & M. A. Moskowitz (1999) Pathobiology of ischaemic stroke: an integrated view. Trends Neurosci, 22, 391–7.

Fisher, M., G. Feuerstein, D. W. Howells, P. D. Hurn, T. A. Kent, S. I. Savitz & E. H. Lo (2009) Update of the Stroke Therapy Academic Industry Roundtable Preclinical Recommendations. Stroke, 40, 2244–2250.

Fitch, M. T., C. Doller, C. K. Combs, G. E. Landreth & J. Silver (1999) Cellular and molecular mechanisms of glial scarring and progressive cavitation: in vivo and in vitro analysis of inflammation-induced secondary injury after CNS trauma. J Neurosci, 19, 8182–98.

Ford, G., Z. Xu, A. Gates, J. Jiang & B. D. Ford (2006) Expression Analysis Systematic Explorer (EASE) analysis reveals differential gene expression in permanent and transient focal stroke rat models. Brain Res, 1071, 226–36.

Giulian, D., M. Corpuz, S. Chapman, M. Mansouri & C. Robertson (1993a) Reactive mononuclear phagocytes release neurotoxins after ischemic and traumatic injury to the central nervous system. J Neurosci Res, 36, 681–93.

Giulian, D., K. Vaca & M. Corpuz (1993b) Brain glia release factors with opposing actions upon neuronal survival. J Neurosci, 13, 29–37.

Guglielmotto, M., M. Aragno, R. Autelli, L. Giliberto, E. Novo, S. Colombatto, O. Danni, M. Parola, M. A. Smith, G. Perry, E. Tamagno & M. Tabaton (2009) The up-regulation of BACE1 mediated by hypoxia and ischemic injury: role of oxidative stress and HIF1alpha. J Neurochem, 108, 1045–56.

Hallenbeck, J. M. (1996) Significance of the inflammatory response in brain ischemia. Acta Neurochir Suppl, 66, 27–31.

Hossmann, K. A. (2006) Pathophysiology and therapy of experimental stroke. Cell Mol Neurobiol, 26, 1057–83.

Hyman, M. C., D. Petrovic-Djergovic, S. H. Visovatti, H. Liao, S. Yanamadala, D. Bouïs, E. J. Su, D. A. Lawrence, M. J. Broekman, A. J. Marcus & D. J. Pinsky (2009) Self-regulation of inflammatory cell trafficking in mice by the leukocyte surface apyrase CD39. J Clin Invest, 119, 1136–49.

Iadecola, C. & M. Alexander (2001) Cerebral ischemia and inflammation. Curr Opin Neurol, 14, 89–94.

Iadecola, C. & J. Anrather (2011) The immunology of stroke: from mechanisms to translation. Nat Med, 17, 796–808.

Ito, D., K. Tanaka, S. Suzuki, T. Dembo & Y. Fukuuchi (2001) Enhanced expression of Iba1, ionized calcium-binding adapter molecule 1, after transient focal cerebral ischemia in rat brain. Stroke, 32, 1208–15.

Jayaraj, R. L., S. Azimullah, R. Beiram, F. Y. Jalal & G. A. Rosenberg (2019) Neuroinflammation: friend and foe for ischemic stroke. J Neuroinflammation, 16, 142.

Jeong, H. K., K. Ji, K. Min & E. H. Joe (2013) Brain inflammation and microglia: facts and misconceptions. Exp Neurobiol, 22, 59–67.

Jiang, L., H. Mu, F. Xu, D. Xie, W. Su, J. Xu, Z. Sun, S. Liu, J. Luo, Y. Shi, R. K. Leak, L. R. Wechsler, J. Chen & X. Hu (2020) Transcriptomic and functional studies reveal undermined chemotactic and angiostimulatory properties of aged microglia during stroke recovery. J Cereb Blood Flow Metab, 40, S81–S97.

Kawabori, M. & M. A. Yenari (2015) The role of the microglia in acute CNS injury. Metab Brain Dis, 30, 381–92.

Kim, J. Y., J. Park, J. Y. Chang, S. H. Kim & J. E. Lee (2016) Inflammation after Ischemic Stroke: The Role of Leukocytes and Glial Cells. Exp Neurobiol, 25, 241–251.

Kindy, M. S., A. N. Bhat & N. R. Bhat (1992) Transient ischemia stimulates glial fibrillary acid protein and vimentin gene expression in the gerbil neocortex, striatum and hippocampus. Brain Res Mol Brain Res, 13, 199–206.

Koistinaho, M. & J. Koistinaho (2005) Interactions between Alzheimer’s disease and cerebral ischemia--focus on inflammation. Brain Res Brain Res Rev, 48, 240–50.

Li, L., X. Zhang, D. Yang, G. Luo, S. Chen & W. Le (2009) Hypoxia increases Abeta generation by altering beta-and gamma-cleavage of APP. Neurobiol Aging, 30, 1091–8.

Li, S. L., J. J. Overman, D. Katsman, S. V. Kozlov, C. J. Donnelly, J. L. Twiss, R. J. Giger, G. Coppola, D. H. Geschwind & S. T. Carmichael (2010) An age-related sprouting transcriptome provides molecular control of axonal sprouting after stroke. Nature Neuroscience, 13, 1496–U82.

Lipton, P. (1999) Ischemic cell death in brain neurons. Physiol Rev, 79, 1431–568.

Liu, Y., X. Xue, H. Zhang, X. Che, J. Luo, P. Wang, J. Xu, Z. Xing, L. Yuan, X. Fu, D. Su, S. Sun, C. Wu & J. Yang (2019) Neuronal-targeted TFEB rescues dysfunction of the autophagy-lysosomal pathway and alleviates ischemic injury in permanent cerebral ischemia. Autophagy, 15, 493–509.

Matsumoto, H., Y. Kumon, H. Watanabe, T. Ohnishi, M. Shudou, C. Ii, H. Takahashi, Y. Imai & J. Tanaka (2007) Antibodies to CD11b, CD68, and lectin label neutrophils rather than microglia in traumatic and ischemic brain lesions. J Neurosci Res, 85, 994–1009.

Noll, J. M., Y. Li, T. J. Distel, G. D. Ford & B. D. Ford (2019) Neuroprotection by Exogenous and Endogenous Neuregulin-1 in Mouse Models of Focal Ischemic Stroke. J Mol Neurosci, 69, 333–342.

Nowicka, D., K. Rogozinska, M. Aleksy, O. W. Witte & J. Skangiel-Kramska (2008) Spatiotemporal dynamics of astroglial and microglial responses after photothrombotic stroke in the rat brain. Acta Neurobiol Exp (Wars), 68, 155–68.

Pekny, M. & M. Nilsson (2005) Astrocyte activation and reactive gliosis. Glia, 50, 427–34.

Price, C. J., D. Wang, D. K. Menon, J. V. Guadagno, M. Cleij, T. Fryer, F. Aigbirhio, J. C. Baron & E. A. Warburton (2006) Intrinsic activated microglia map to the peri-infarct zone in the subacute phase of ischemic stroke. Stroke, 37, 1749–53.

Rodriguez-Mercado, R., G. D. Ford, Z. F. Xu, E. N. Kraiselburd, M. I. Martinez, V. A. Eterovic, E. Colon, I. V. Rodriguez, P. Portilla, P. A. Ferchmin, L. Gierbolini, M. Rodriguez-Carrasquillo, M. D. Powell, J. V. K. Pulliam, C. O. McCraw, A. Gates & B. D. Ford (2012) Acute Neuronal Injury and Blood Genomic Profiles in a Nonhuman Primate Model for Ischemic Stroke. Comparative Medicine, 62, 427–438.

Sanchez-Bezanilla, S., R. J. Hood, L. E. Collins-Praino, R. J. Turner, F. R. Walker, M. Nilsson & L. K. Ong (2021) More than motor impairment: A spatiotemporal analysis of cognitive impairment and associated neuropathological changes following cortical photothrombotic stroke. J Cereb Blood Flow Metab, 271678×211005877.

Savchenko, V. L., J. A. McKanna, I. R. Nikonenko & G. G. Skibo (2000) Microglia and astrocytes in the adult rat brain: comparative immunocytochemical analysis demonstrates the efficacy of lipocortin 1 immunoreactivity. Neuroscience, 96, 195–203.

Simmons, L. J., M. C. Surles-Zeigler, Y. G. Li, G. D. Ford, G. D. Newman & B. D. Ford (2016) Regulation of inflammatory responses by neuregulin-1 in brain ischemia and microglial cells in vitro involves the NF-kappa B pathway. Journal of Neuroinflammation, 13.

Stoll, G., S. Jander & M. Schroeter (1998) Inflammation and glial responses in ischemic brain lesions. Prog Neurobiol, 56, 149–71.

Sun, X., G. He, H. Qing, W. Zhou, F. Dobie, F. Cai, M. Staufenbiel, L. E. Huang & W. Song (2006) Hypoxia facilitates Alzheimer’s disease pathogenesis by up-regulating BACE1 gene expression. Proc Natl Acad Sci U S A, 103, 18727–32.

Surles-Zeigler, M. C., Y. G. Li, T. J. Distel, H. Omotayo, S. K. Ge & B. D. Ford (2018) Transcriptomic analysis of neuregulin-1 regulated genes following ischemic stroke by computational identification of promoter binding sites: A role for the ETS-1 transcription factor. Plos One, 13.

Wang, R., Y. W. Zhang, X. Zhang, R. Liu, S. Hong, K. Xia, J. Xia, Z. Zhang & H. Xu (2006) Transcriptional regulation of APH-1A and increased gamma-secretase cleavage of APP and Notch by HIF-1 and hypoxia. FASEB J, 20, 1275–7.

Wang, S. L., Y. G. Li, R. Paudyal, B. D. Ford & X. D. Zhang (2015) Spatio-temporal assessment of the neuroprotective effects of neuregulin-1 on ischemic stroke lesions using MRI. Journal of the Neurological Sciences, 357, 28–34.

Xu, Z., D. R. Croslan, A. E. Harris, G. D. Ford & B. D. Ford (2006) Extended therapeutic window and functional recovery after intraarterial administration of neuregulin-1 after focal ischemic stroke. J Cereb Blood Flow Metab, 26, 527–35.

Xu, Z., G. D. Ford, D. R. Croslan, J. Jiang, A. Gates, R. Allen & B. D. Ford (2005) Neuroprotection by neuregulin-1 following focal stroke is associated with the attenuation of ischemia-induced pro-inflammatory and stress gene expression. Neurobiol Dis, 19, 461–70.

Xu, Z., J. Jiang, G. Ford & B. D. Ford (2004) Neuregulin-1 is neuroprotective and attenuates inflammatory responses induced by ischemic stroke. Biochem Biophys Res Commun, 322, 440–6.

Zheng, G. Q., X. M. Wang, Y. Wang & X. T. Wang (2010) Tau as a potential novel therapeutic target in ischemic stroke. J Cell Biochem, 109, 26–9.

Zhong, J., A. Chan, L. Morad, H. I. Kornblum, G. Fan & S. T. Carmichael (2010) Hydrogel matrix to support stem cell survival after brain transplantation in stroke. Neurorehabilitation and Neural Repair, 24, 636–644.

Zou, X., X. J. Yang, Y. M. Gan, D. L. Liu, C. Chen, W. Duan & J. R. Du (2021) Neuroprotective Effect of Phthalide Derivative CD21 against Ischemic Brain Injury:Involvement of MSR1 Mediated DAMP peroxiredoxin1 Clearance and TLR4 Signaling Inhibition. J Neuroimmune Pharmacol, 16, 306–317.

